# *µ*bialSim: constraint-based dynamic simulation of complex microbiomes

**DOI:** 10.1101/716126

**Authors:** Denny Popp, Florian Centler

## Abstract

Microbial communities are pervasive in the natural environment, associated with many animal hosts, and of increasing importance in biotechnological applications. The complexity of these microbial systems makes the underlying mechanisms driving their dynamics difficult to identify. While experimental meta-OMICS techniques are routinely applied to record the inventory and activity of microbiomes over time, it remains difficult to obtain quantitative predictions based on such data. Mechanistic, quantitative mathematical modeling approaches hold the promise to both provide predictive power and shed light on cause-effect relationships driving these dynamic systems. We introduce *µ*bialSim (pronounced “microbialsim”), a dynamic Flux-Balance-Analysis-based (dFBA) numerical simulator which is able to predict the time course in terms of composition and activity of microbiomes containing 100s of species in batch or chemostat mode. Activity of individual species is simulated by using separate FBA models which have access to a common pool of compounds, allowing for metabolite exchange. A novel augmented forward Euler method ensures numerically accuracy by temporarily reducing the time step size when compound concentrations decrease rapidly due to high compound affinities and/or the presence of many consuming species. We present three exemplary applications of *µ*bialSim: a batch culture of a hydrogenotrophic archaeon, a syntrophic methanogenic biculture, and a 773-species human gut microbiome which exhibits a complex and dynamic pattern of metabolite exchange.

Focussing on metabolite exchange as the main interaction type, *µ*bialSim allows for the mechanistic simulation of microbiomes at their natural complexity. Simulated trajectories can be used to contextualize experimental meta-OMICS data, and hypotheses on cause-effect relationships driving community dynamics can be derived based on scenario simulations.

*µ*bialSim is implemented in Matlab and relies on the COBRA Toolbox or CellNetAnalyzer for FBA calculations. The source code is available under the GNU General Public License v3.0 at https://git.ufz.de/UMBSysBio/microbialsim.

## Introduction

Microbial communities are ubiquitous in nature, thriving in diverse habitats ranging from the deep subsurface [1] over digestive tracts of higher animals [2] to the upper troposphere [3]. They are self-organizing entities which both modulate the environment they are embedded in, as well as their own constituents in terms of abundance of individual member populations. Typical natural and engineered microbiomes engage in numerous metabolic and non-metabolic interactions and contain a large fraction of not-yet cultured species. The resulting complexity makes microbiomes notoriously difficult to study. Meta-OMICS techniques help to uncover the metabolic potential and current activity of microbiomes. However, most analyses based on such data remains observational in nature and cannot be used to derive quantitative predictions. The mathematical modeling of microbiomes holds the promise to move from observation to a more quantitative understanding of microbiome dynamics and underlying mechanisms [4–7].

Focusing on metabolic interaction, a number of dynamic community modeling approaches have been proposed in which activity of individual species is modeled using constraint-based techniques based on genome-scale metabolic network reconstructions [8]. Some of these approaches require the definition of a secondary community objective in addition to the standard growth maximization objective for individual species (e.g., d-OptCom, [9]), a priority list of objectives (DFBAlab [10]), or a pre-allocation of compounds to competing species [11]. Other models additionally allow for parameter calibration (MCM [12]), or for the inclusion of space either simulating populations (COMETS [13], MetaFlux [14]) or individual microbial cells following a rule-based approach (BacArena, [15]). With the exception of the last approach, typically only microbiomes of few species have been considered in simulations yet. In order to be able to mirror the diversity of natural microbiomes, we developed *µ*bialSim. Our simulator is based on the dynamic Flux-Balance-Analysis approach and does not require the definition of any additional objectives or the pre-allocation of compounds. It allows for the simulation of well-mixed microbiomes of high diversity under batch and chemostat conditions with high numerical accuracy due to a novel numerical integration scheme.

## Design and Implementation

### Overview

In order to simulate the fate and metabolic activity of a microbial community we follow the compartmentalized approach in which activity and growth of individual species is modeled by separate genome-scale metabolic network models following the Flux-Balance-Analysis approach (FBA, [16]). All species have access to a common set of pool compounds. This allows for competition between species as they try to consume the same pool compound and cross-feeding if one species produces a pool compound another is able to use for growth. Instead of restricting analysis to steady state dynamics for which the community composition must be defined as a model input (e.g., [17,18]), we follow the dynamic FBA approach [19] in order to be able to simulate dynamic shifts in microbiomes as a consequence of the system’s dynamics. In this approach, the steady-state assumption underlying FBA is assumed to hold true for the duration of the numerical integration step. FBA-computed growth and compound exchange rates are then used to update the state variables of the model which encompass microbial biomass and pool compound concentrations. *µ*bialSim is implemented as Matlab code and relies on either the COBRA Toolbox [20] or CellNetAnalyzer [21] for performing FBA computations. This allows for the easy incorporation of FBA models prepared with either softwares in a community model. Space is neglected in the model, hence assuming a well-mixed environment similar to a well-stirred bioreactor. Both batch and chemostat operation can be simulated. Both compounds and microbial populations can be defined to be part of the bioreactor inflow.

### Mathematical description

The system state is given by (*C*, *X*), with *C* = (*C*_1_,…,*C*_*m*_) referring to the concentrations (in mM) of *m* pool compounds present in the bioreactor and *X* = (*X*_1_,…,*X*_*n*_) referring to the abundance (in gDW/L) of *n* microbial populations. For each of these populations, the exchange reactions in their metabolic network model which describe the transport of a metabolite across the cell membrane need to be identified. Not all of these reactions need to be coupled to pool compounds. For example metabolites assumed not to be growth-limiting can be ignored. With *k* the number of coupled exchange reactions for species *j*, *coupReac*^*j*^ = (*r*_1_,…,*r*_*k*_) records the reaction IDs of the respective exchange reactions, *coupComp*^*j*^=(*idx*_*i*_,…, *idx*_*k*_) the indices of the corresponding compounds in *C*, *coupSense*^*j*^=(*s*_*i*_,…, *s*_*k*_) the directionality of the exchange reaction with the reaction proceeding in the forward direction indicating metabolite excretion for *s* = 1 and metabolite uptake for *s* = −1, *coupVmax*^*j*^ the maximal uptake fluxes, and *coupKs*^*j*^ the corresponding Monod constants (see below). The dynamics of the system is then given by two sets of ordinary differential equations. Microbial dynamics for species *j* is given by

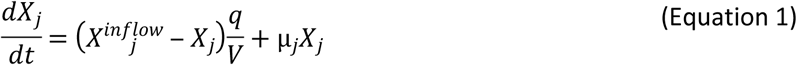

with microbial concentration in the inflow 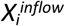 (gDW/L), flow rate q (L/h), bioreactor volume V (L), and specific growth rate *µ*_*i*_ (1/h). The dynamics of pool compound *i* in the bioreactor is given by

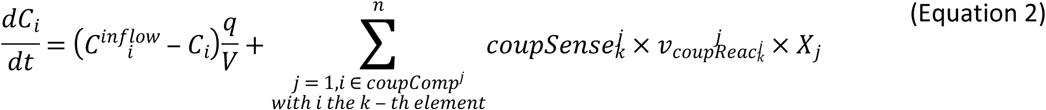

with inflow concentration 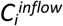 (mM) and flux of the exchange reaction 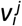 (mmol/gDW/h) which is the *i*-th reaction of the *j*-th species.

The specific growth rates *µ* and exchange fluxes *v* are derived by solving individual FBA problems for all species individually. For this purpose, current compound concentrations in the bioreactor need to be translated to maximal allowable uptake rates. This is commonly done by assuming Monod-type kinetics. For the *i*-th exchange reaction of species *j* which is coupled to pool compound *coupComp*^*j*^_*i*_, the current maximal uptake rate is given by

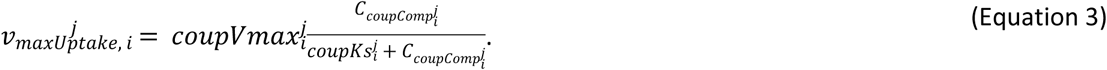

### Numerical integration scheme

While *µ*bialSim can make use of Matlab solvers for numerically integrating Equations 1–2 (options `solverPars.solverType` and `solverPars.solver`), the computational costs quickly becomes prohibitive for more complex microbial communities. Instead, we have implemented a novel augmented forward Euler method in *µ*bialSim. The forward Euler method uses the system state at time *t*, evaluates Equations 1–2 and uses computed rates to derive the system state at time *t* + Δ*t*, with Δ*t* being the integration step size:

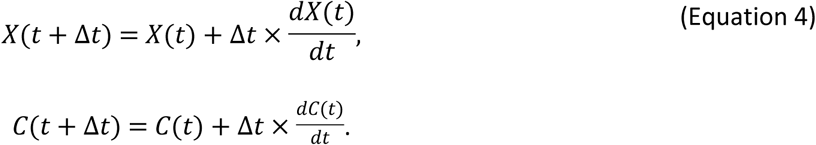

For syntrophic interactions such as in syntrophic propionate degradation (see Example 2), a compound produced by one species (here: hydrogen), needs to be quickly consumed by the syntrophic partner (here: a methanogenic archaeon) as propionate degradation is thermodynamically only feasible for low hydrogen concentrations. This means that typically, the partner features an effective uptake of the compound with a small *K*_*s*_ value in Equation 3. As consumption can become much faster than production, a very negative rate for hydrogen may result in Equation 2. This can lead to the computation of negative concentrations during an integration step (Equation 4). Similarly, this can also be caused by many species competing for a highly attractive compound. Simply setting negative values to zero in each integration step induces a numerical error. Instead, choosing a smaller integration step size can solve this problem, but might significantly prolong simulation time. Hence, in *µ*bialSim the integration step size is reduced only temporarily whenever this situation occurs in order to avoid numerical error at an affordable increase in computational cost. The time step size is reduced in such a way that the concentration of compound *o* at the next time step is close to its steady-state concentration under the assumption that the production process remains constant. We first identify all species which are either producing or consuming compound *o*. We then compute the current total production rate *p* and the current total uptake rate *u* for the compound by summing across the identified species. Additionally, let *f* describe the current rate of concentration change for compound *o* due to a prescribed flow if a chemostat is simulated. The steady-state condition is then given by *p* = *u* -*f*. Treating *p* as fixed, we find that the right-hand side of this equation depends on the compound concentration *C*_*o*_ when combining Equations 2 and 3:

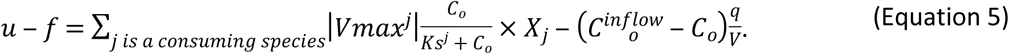

Under the assumption that compound *o* is the growth-limiting factor for the second species (i.e., the maximal uptake rate is indeed realized) and that growth remains viable for smaller concentrations, the steady-state concentration *C*^*^_*o*_ for compound *o* can be found by reducing concentration *C*_*o*_ in Equation 5 until *p* = *u*(*C*^*^_*o*_)-*f*(*C*^*^_*o*_). The time step size Δ*t* which leads *C*_o_ (*t* + Δ*t*) to be evaluated to *C*^*^_*o*_ can then be computed with the help of Equation 4 to:

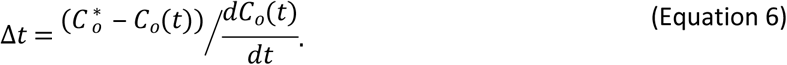

If for more than one chemical compound negative concentrations were calculated using the default time step size, for each of these compounds the described scheme is applied and ultimately the smallest time step size used. We note that reducing the time step size does not require the recomputation as FBA problems, as only Δ*t* changes in Equation 4. For the next time step, the default time step size is restored. Compounds which required the reduction of the time step size are flagged as strongly consumed compounds, as their consumption rate surpassed their production rate. In order to avoid oscillatory behavior for these compounds, *µ*bialSim allows to additionally restrict the time step size in subsequent iteration steps such that the concentration change of these compounds does not surpass a given threshold (parameter `solverPars.maxDeviation`). If negative biomass concentrations occur, the time step size is reduced such that the biomass concentration is at most reduced by a factor of two. The flowchart in Fig 1 depicts the complete algorithmic logic of the augmented forward Euler method implemented in *µ*bialSim.

**Fig 1.**
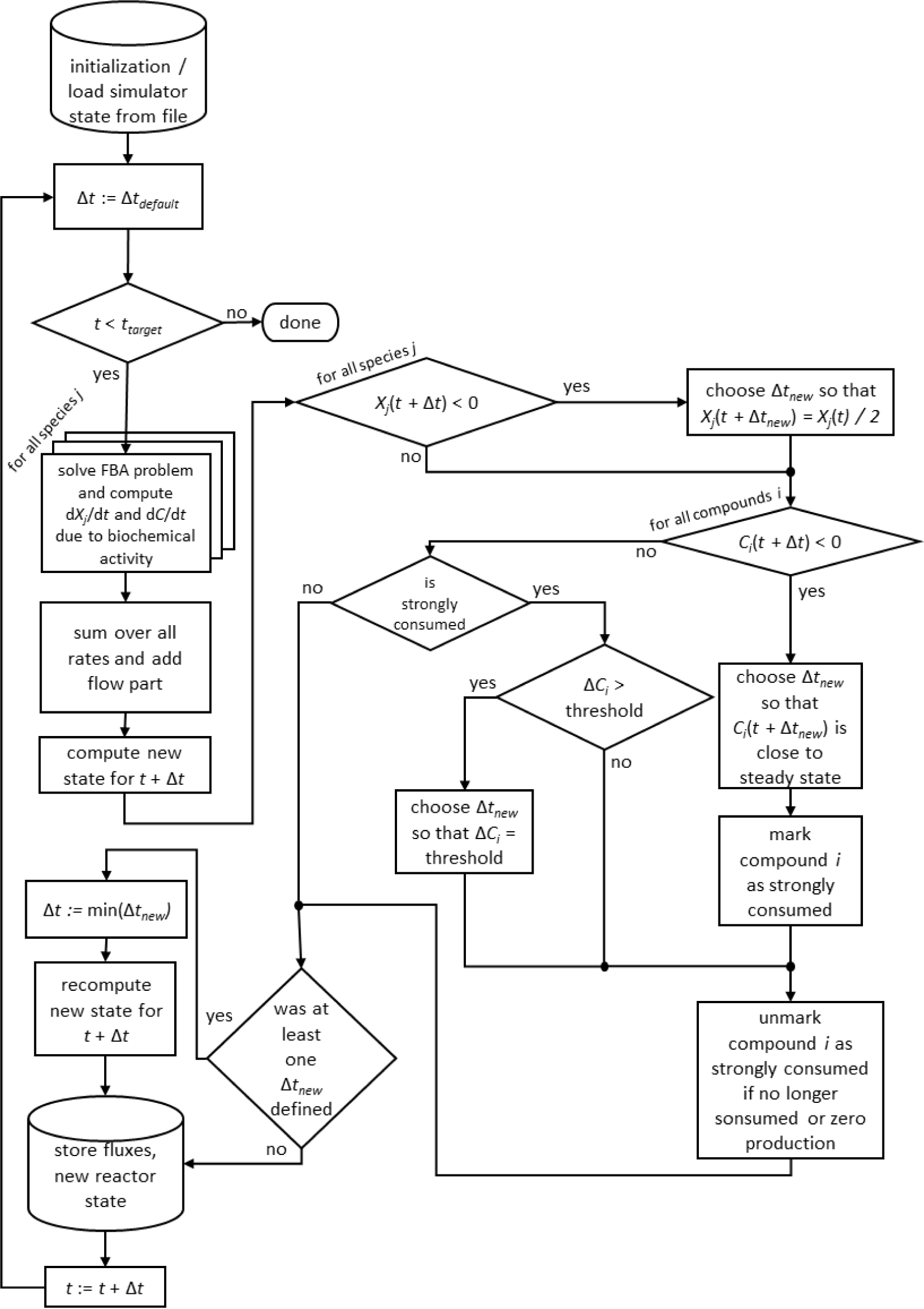
The augmented forward Euler scheme implemented in *µ*bialSim. In each numerical integration step, first the FBA solutions are computed for all member species of the simulated microbiome. The new system state is then computed using obtained rates and the default time step size. If negative concentration values for biomasses or compounds occur, the time step size is reduced as required.

### Features

FBA computations can have non-unique solutions such that different flux distributions lead to the same maximal growth rate. In dFBA simulations, this can cause discontinuities in intracellular fluxes over time. To avoid this, *µ*bialSim implements two features which can individually or in tandem be activated. The first feature is a secondary optimization step which seeks to realize the optimal growth rate as determined by the initial FBA computation, but with minimal fluxes, known as parsimonious FBA [22]. The second feature tries to realize the optimal growth rate by a flux distribution that most resembles the flux distribution which was active in the last integration step, a methodology similar to the minimization of metabolic adjustment approach (MOMA, [23]) which has been applied in the context of dFBA before [24]. Simulation results can be stored at each integration step in individual files or in a single result file at the end of the simulation. The former feature (parameter `solverPars.recording`) is helpful for complex simulations as simulated data is not lost in case of unforeseen server downtimes or other computational calamities. A subsequent simulator run can use the saved data to initialize the simulator and continue the interrupted simulation run (parameter `solverPars.readInitialStateFrom`).

As loading SBML files and preparing the corresponding data structures can take a while for complex microbiomes, the data structures of the loaded models can be saved as a single file and be used in subsequent simulation runs to speed up initialization (parameter `solverPars.saveLoadedModelToFile`).

Once the simulation is done, *µ*bialSim computes the overall activity during the simulation for all exchange fluxes of all species (including both exchange reactions which were coupled to pool compounds and those which were not) if desired (parameter `solverPars.doMassBalance`). This indicates the total compound turnover per species in terms of compound production minus consumption (in mM), and the resulting increase in biomass concentration (in gDW/L). Additionally, three figures to visualize the simulation result are automatically generated. The first figure gives a quick overview over the temporal evolution of all microbial biomass concentrations and all pool compound concentrations over time. In the second figure, all biomass concentrations are plotted in one panel as an offset to the initial biomass concentration, to make dynamics easy to inspect for species having very different initial biomass concentrations, and individual panels for each pool compound. The third figure contains two panels for each microbial species and shows the evolution of coupled exchange reactions, and exchange reactions which were not coupled. Only non-zero exchange fluxes are shown.

### Setting up and running a microbiome simulation

The bioreactor and its operational parameters are defined in the function `reactorDefinition_*.m`. Here, the reactor volume, flow rate, and the list of pool compounds is defined. Additionally, initial concentrations for compounds and biomasses are specified, as well as their concentration in the inflow in case a chemostat is to be simulated.

Loading a FBA model of an individual species of the microbiome to be simulated is recommended to be done in two steps. First, the model is loaded by using the appropriate commands of either the COBRA Toolbox or CellNetAnalyzer in the Matlab function `prepareFBAmodel_*.m`. After loading, if necessary, general constraints on particular reactions can be set, for example to implement a particular scenario. Next, the reaction IDs of the biomass reaction and the non-growth associated maintenance reaction (NGAM) need to be specified. Reaction IDs refer to their running order in the SBML file (or corresponding CellNetAnalyzer data structure). Furthermore, all exchange reactions need to be identified by their IDs and their directionality, that means whether a positive flux indicates compound secretion (`Sense` = 1) or compound uptake (`Sense` = −1). Finally the subset of exchange reactions are identified, which will be coupled to pool compounds present in the bioreactor in the vector `IDs`. The mapping of coupled reactions to reactor compounds is done in the vector `reactorCompoundIDs` of length *k*, with *k* indicating the number of coupled reactions. The entry at the *i*-th position specifies for the *i*-th coupled reaction, as defined before in the vector `IDs`, the index of the reactor compound (referring to vector `reactor.compounds`) to which the exchange reaction is coupled. After this general setup of the FBA model, model parameters are defined in the second step in the function `parametrizeFBAmodel_*.m`. Here, the values for NGAM, and *v*_*max*_ and *K*_*S*_ to define uptake kinetics for all coupled compounds are set.

Finally, the target simulation time, default time step size and other options (see Features) and numerical accuracy parameters are set in the main simulator file `microbialSimMain.m`.

## Results

We present three exemplary applications of *µ*bialSim simulating batch growth of a monoclonal hydrogenotrophic culture, a syntrophic biculture transforming propionate to methane, and a 773 species human gut microbiome. In all examples, a bioreactor volume of 1 L and a default time step size of Δ*t*_*default*_ = 0.002 h was chosen. The simulation end time was set to *t*_*target*_ = 1 h for the mono- and binary culture, and to 0.3 h for the human microbiome example. All simulations were run in Matlab R2018a on an Intel^®^ Xeon^®^ CPU E5-4620 v2@2.6GHz with 32 cores. Up to 64GB of RAM were required to simulate the 773 species microbiome.

### Batch culture of *Methanococcus maripaludis*

A batch culture of the hydrogenotrophic methanogen *M. maripaludis* was simulated using an established genome-scale FBA model [25]. The archaeon transforms H_2_ and CO_2_ to CH_4_. Excess CO_2_ was provided such that H_2_ was the growth limiting factor. Model parameters and initial conditions are listed in Table 1. Simulation results show an almost linear growth of *M. maripaludis* until *t* = 0.6 h when H_2_ becomes depleted and growth stops (Fig 2). Simulations using Matlab’s ODE solver ode15s and the novel augmented forward Euler method lead to identical results (Fig 2) with comparable simulation times (2.2 minutes for Matlab’s solver and 3.4 minutes for the Euler method).

**Table 1.** Model parameters and initial conditions for Example 1.

| Model parameters for *M. maripaludis* model iMR539 |  |  |  | Initial conditions |  |  |
| --- | --- | --- | --- | --- | --- | --- |
| $\mu$ (1/d) | $v_{max}^a$<br>(mmol/<br>gDW/h) | $K_s$ (mM) | NGAM<br>(mmol<br>ATP/gDW/h) | Biomass<br>(gDW/L) | H<sub>2</sub> (mM) | CO<sub>2</sub> (mM) |
| 2.1 [26] | 189.3 | $4.375 \times 10^{-4}$ [26] | 5.1176 [25] | $1.0 \times 10^{-4}$ | 0.01 | 1.0 |

<sup>a</sup>Was chosen such that the maximal FBA-predicted growth rate matched the specific growth rate $\mu$ reported in first table column.

**Fig 2.**
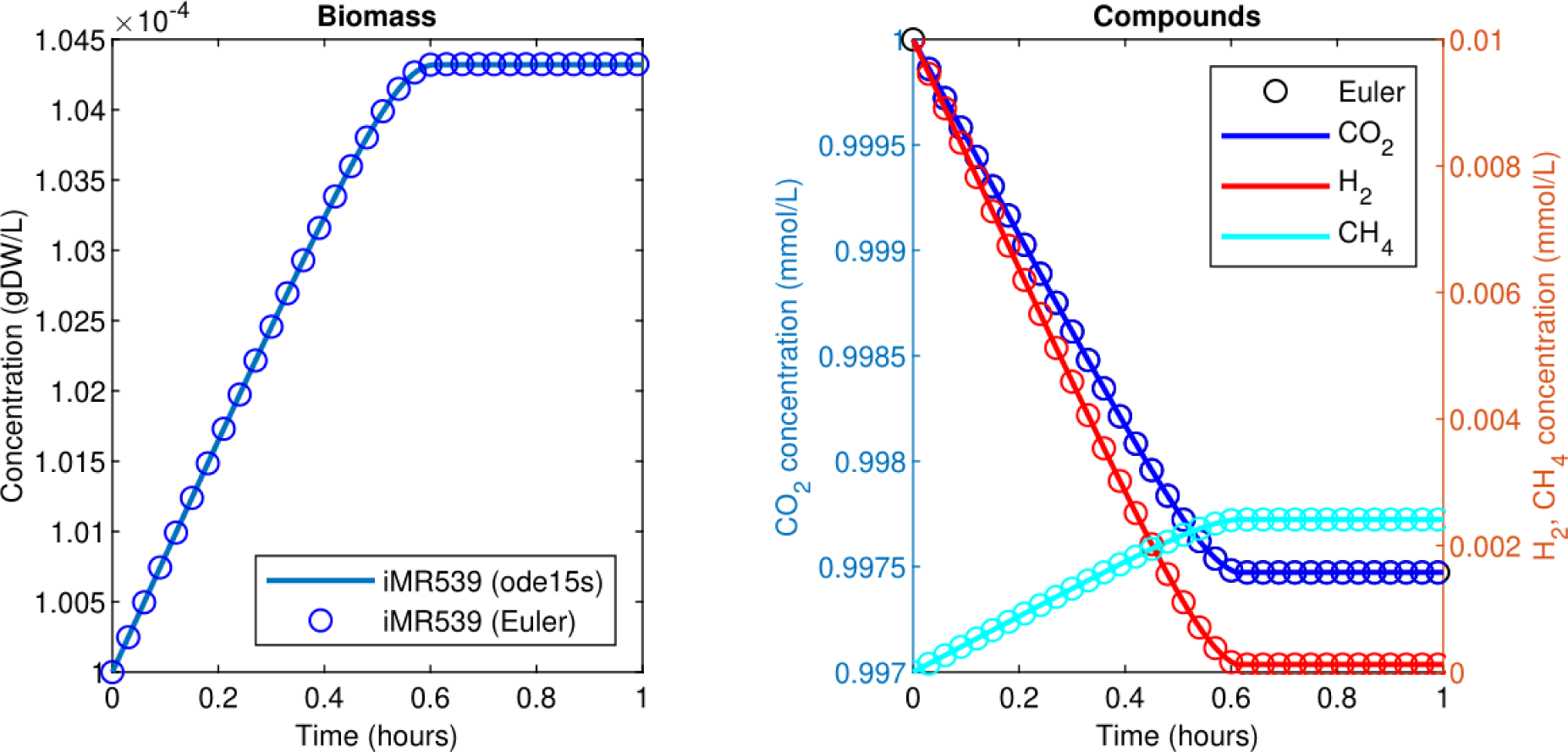
Simulating a hydrogenotrophic batch culture. A *M. maripaludis* population converts H_2_ and CO_2_ to CH_4_ until H_2_ becomes depleted. Both Matlab’s ode15s ODE solver (lines) and *µ*bialSim’s novel augmented forward Euler method (symbols, every 15th data point is plotted) lead to identical results.

### Co-culture of *Syntrophobacter fumaroxidans* and *Methanospirillum hungatei*

The syntrophic conversion of propionate to methane was simulated by using a binary FBA model community of *S. fumaroxidans* and *M. hungatei* which has previously been simulated at steady state [18]. Model parameters are listed in Table 2, choosing an initial relative biomass ratio of 3:4 (*M. hungatei*:*S. fumaroxidans*) as previously [18]. Initial compound concentrations were set to 20 mM for propionate, 0.9561 µM for H_2_ and 8.215 µM for CO_2_ which was considered not to be growth limiting for the methanogen. Being produced by *S. fumaroxidans* and quickly consumed by *M. hungatei*, H_2_ was flagged as a strongly consumed compound in the simulation. The time step size became reduced and reached a minimum just prior to the depletion of H_2_ as growth of *S. fumaroxidans* ceased due to low propionate concentrations at *t* = 0.76 h (Fig 3). Except for H_2_, simulation results agreed well if using Matlab’s ODE solver or the novel numerical Euler scheme. For H_2_, minor fluctuations around the ODE result were apparent when using the Euler scheme (Fig 3). Most notably, the final H_2_ concentration was 0 instead of the ODE predicted (small) concentration of 43.9 pM. Simulation times remained below 30 minutes for both Matlab’s ODE solver (9.7 minutes) and the augmented forward Euler method (22.6 minutes).

**Table 2.**
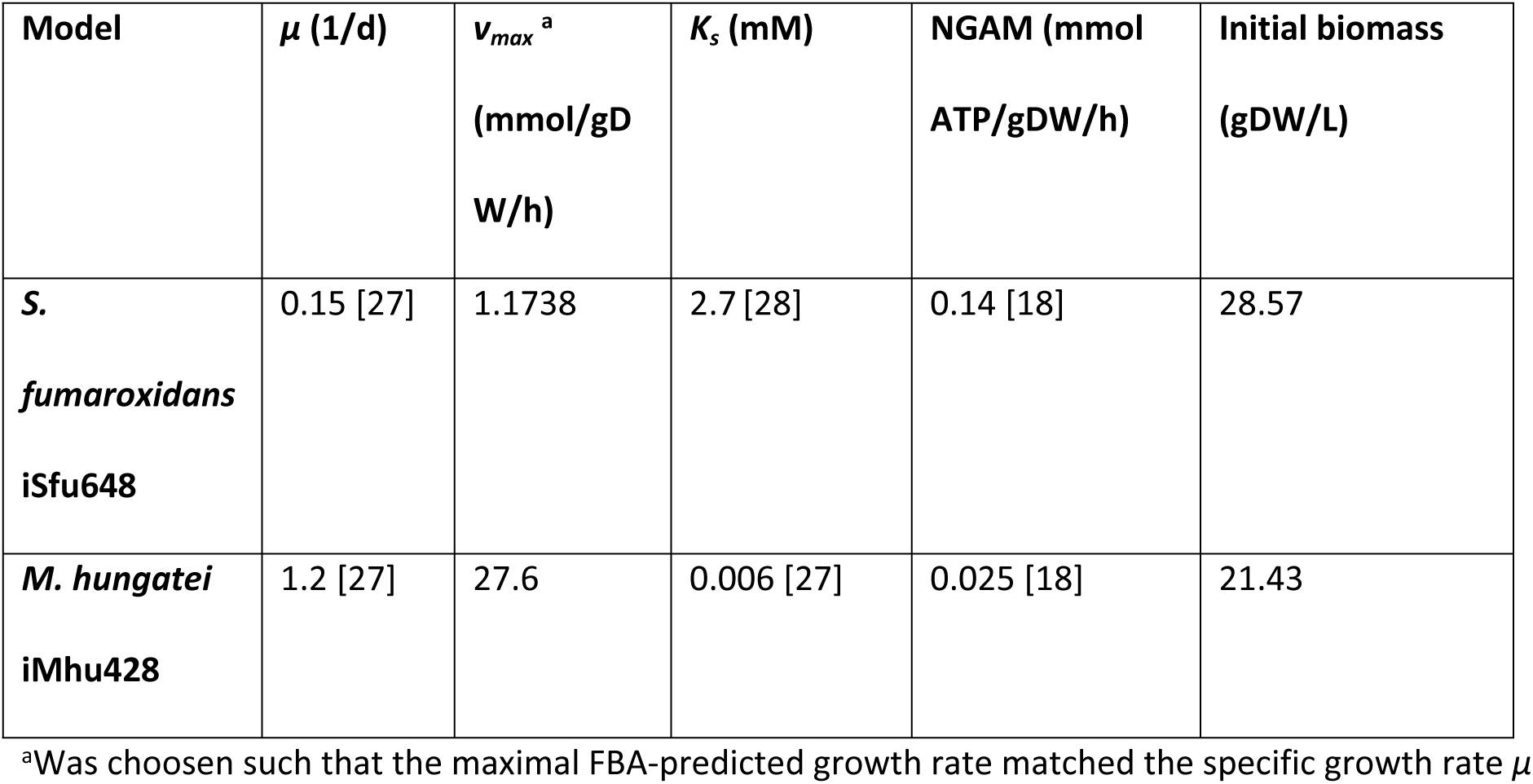
Model parameters and initial biomass concentrations for Example 2.

**Fig 3.**
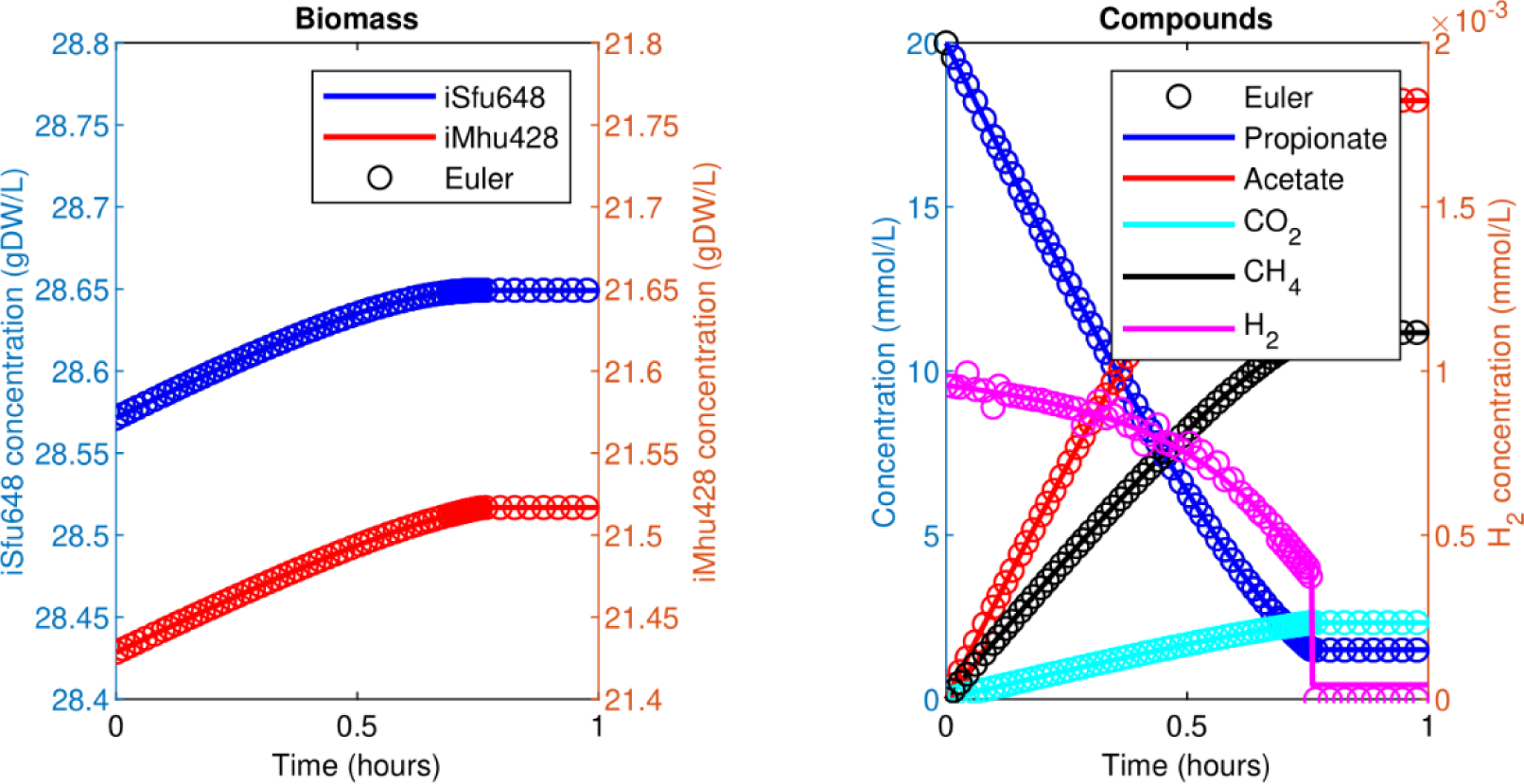
Simulating a binary, syntrophic batch culture. Propionate is utilized by *S. fumaroxidans* and converted to acetate, CO_2_, and H_2_. *M. hungatei* then converts CO_2_ and H_2_ to CH_4_. Both Matlab’s ode15s ODE solver (lines) and *µ*bialSim’s novel augmented forward Euler method (symbols, every 15th data point is plotted) lead to similar results. As H_2_ is faster consumed than produced, the time step size gets frequently reduced, most notably just prior to the depletion of propionate after which growth of both populations ceases.

### Human gut microbiome

To simulate a human gut microbiome, the AGORA model collection (Version 1.01) comprising 773 microbial human gut species was used [29]. Maximal substrate uptake rates (*v*_*max*_) were taken from the individual SBML models, which were configured to mimic a typical western diet [29]. Exchange reactions in individual models were automatically identified by searching for `“EX_”` in reaction names. Pool compounds were automatically configured by considering only those exchange reaction which had at least one flux boundary which was neither zero nor unlimited, resulting in 166 pool metabolites if all 773 models are considered in the simulation. Monod constants for compound uptake were set to 0.01 mM for all pool compounds. Batch growth was simulated by setting initial pool compound concentrations to 1.0 mM for all compounds, and initial biomass concentration to 0.1 gDW/L for all microbial species. Simulation results (requiring 7.2 days of simulation time using the augmented forward Euler method) indicate an initial short period of rapid growth which is followed by a prolonged period of slow growth (Fig. 4).

**Fig 4.**
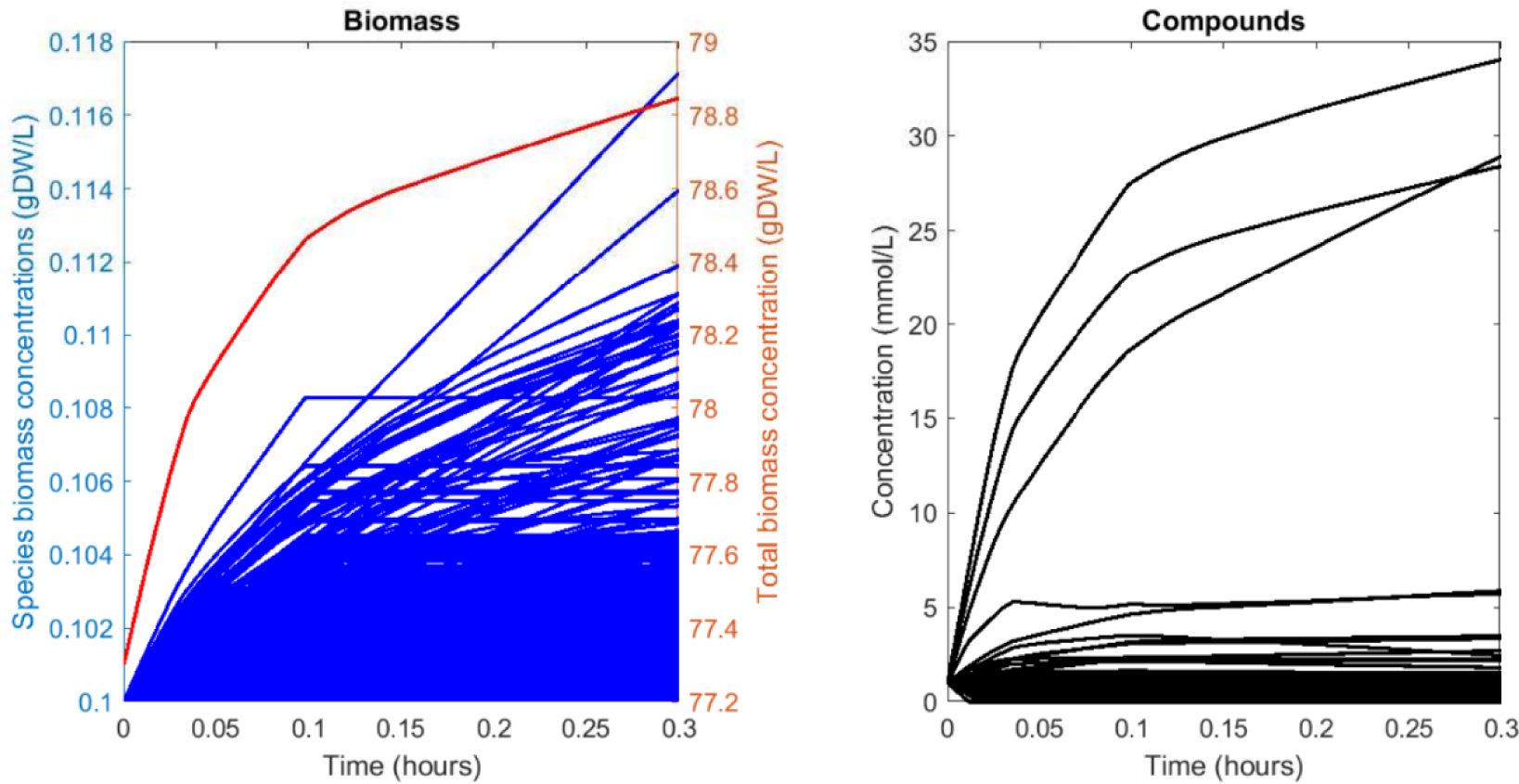
Simulating a 773 species gut microbiome with 166 pool compounds. Total biomass growth slows down as compounds become depleted. Growth for some species ceases early on while others are able to maintain fast growth rates until the end of the simulation.

## Availability and Future Directions

*µ*bialSim is licensed under the GNU General Public License v3.0 and available for download at https://git.ufz.de/UMBSysBio/microbialsim (or `git clone` https://git.ufz.de/UMBSysBio/microbialSim.git). The simulator can make use of Matlab’s support for parallel loop execution (`parfor`, option `solverPar.parallel`) for solving individual FBA problems in one time step. However, the observed speed-up remained far below the expectation of an almost linear speed-up. This is due to the non-persistence of worker processes executing individual loop iterations, requiring the repeated copying of FBA model structures to the workers’ memory in each time step. A future version of *µ*bialSim shall feature persistent workers to better utilize current multicore computing architectures. Besides these technical improvements, non-metabolic interactions as well as chemical activity among pool compounds and non-constant chemostat operating conditions can be implemented in future versions of *µ*bialSim. Furthermore, reactor headspace and corresponding gas exchange processes can be included to ease comparison of simulation results with experimental data.

## Supplemental Material

### Installing *µ*bialSim

The simulator *µ*bialSim is implemented as Matlab code and can be obtained from the UFZ git server at https://git.ufz.de/UMBSysBio/microbialsim or via `git clone` https://git.ufz.de/UMBSysBio/microbialSim.git. *µ*bialSim can be configured to use the COBRA Toolbox or CellNetAnalyzer for performing FBA calculations. The provided examples make use of the former. After installing the COBRA Toolbox, the appropriate path needs to be configured in lines 87ff in the main simulator file `microbialSimMain.m`.

### Simulation output

Two files are generated at the end of the simulation with a date and time stamp in the filename indicating the start of the simulation. Both files hold Matlab data structures. The file `“*_restartInit.mat”` records the final state of the simulator and can be used as the initial conditions to continue the simulation in a subsequent run of *µ*bialSim. The other file holds the simulated trajectory in the Matlab structure `trajectory`. The fields `time, compounds, biomass`, and `mu` hold the time, compound concentrations, biomass concentrations, and specific growth rates for each integration step. The field `FBA` stores data for each FBA model, including the temporal dynamics of all metabolic fluxes, and the mass balance for all exchange reactions.

### Running the examples

#### Example 1: methanogenic monoculture

The first example in which batch-culture growth of a single hydrogenotrophic species (*Methanococcus maripaludis*) is simulated can be run with the command `microbialSimMain(1)`. Once the simulation is finished, the trajectory is automatically visualized in three Matlab figures (Fig S1).

**Fig S1.**
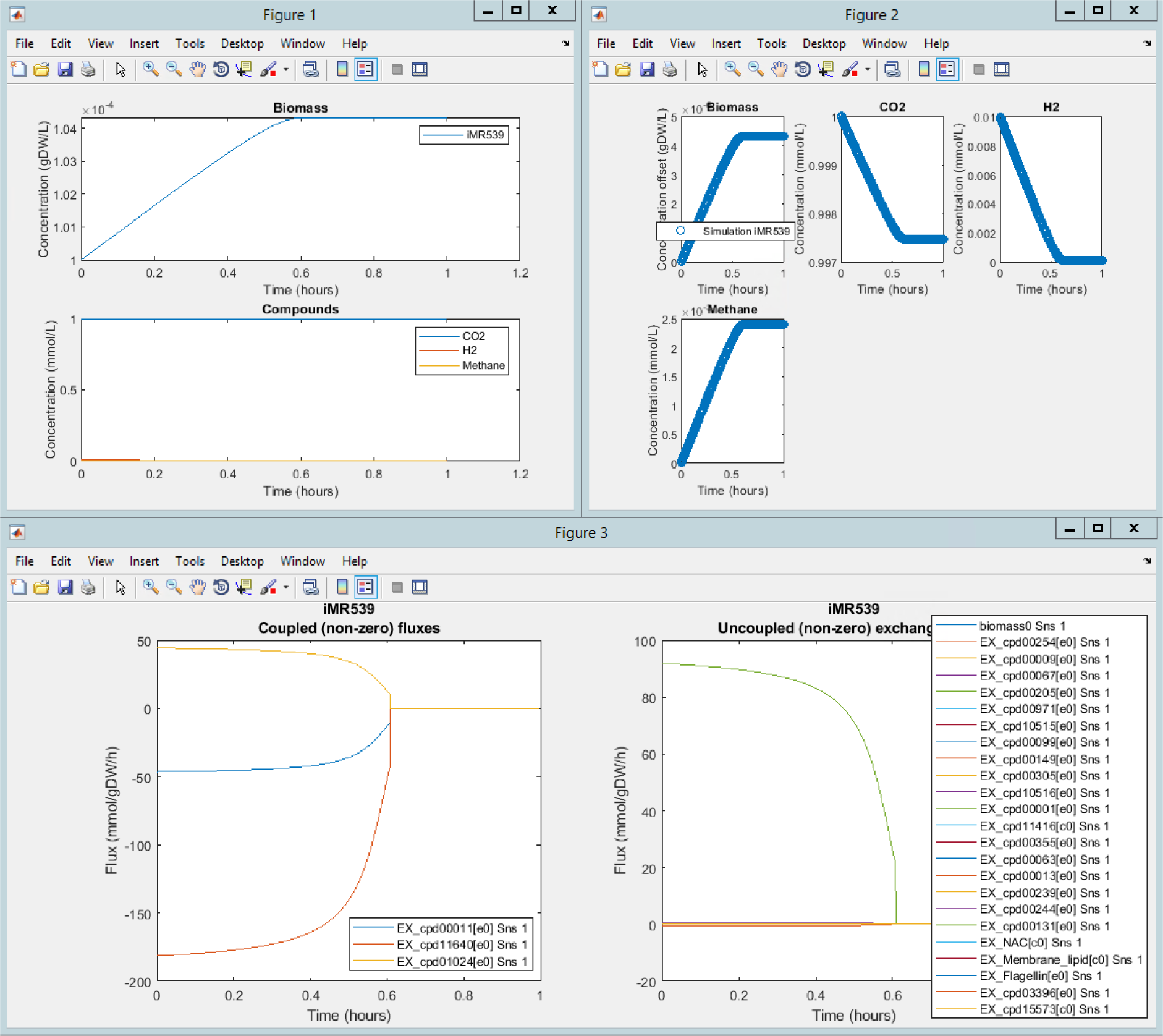
Automatically generated figure for Example 1. Simulation trajectory showing all microbiome member species and reactor compounds (Figure 1). Plot of biomass concentration as an offset to the initial concentration for all microbiome member species and individual plots for all reactor compounds (Figure 2). Plotting non-zero exchange fluxes over time which are coupled to reactor compounds (left), or not (right) for all microbiome member species (Figure 3).

#### Example 2: binary syntrophic community

Batch-culture growth of binary methanogenic community is started by `microbialSimMain(2)`.

**Fig S2.**
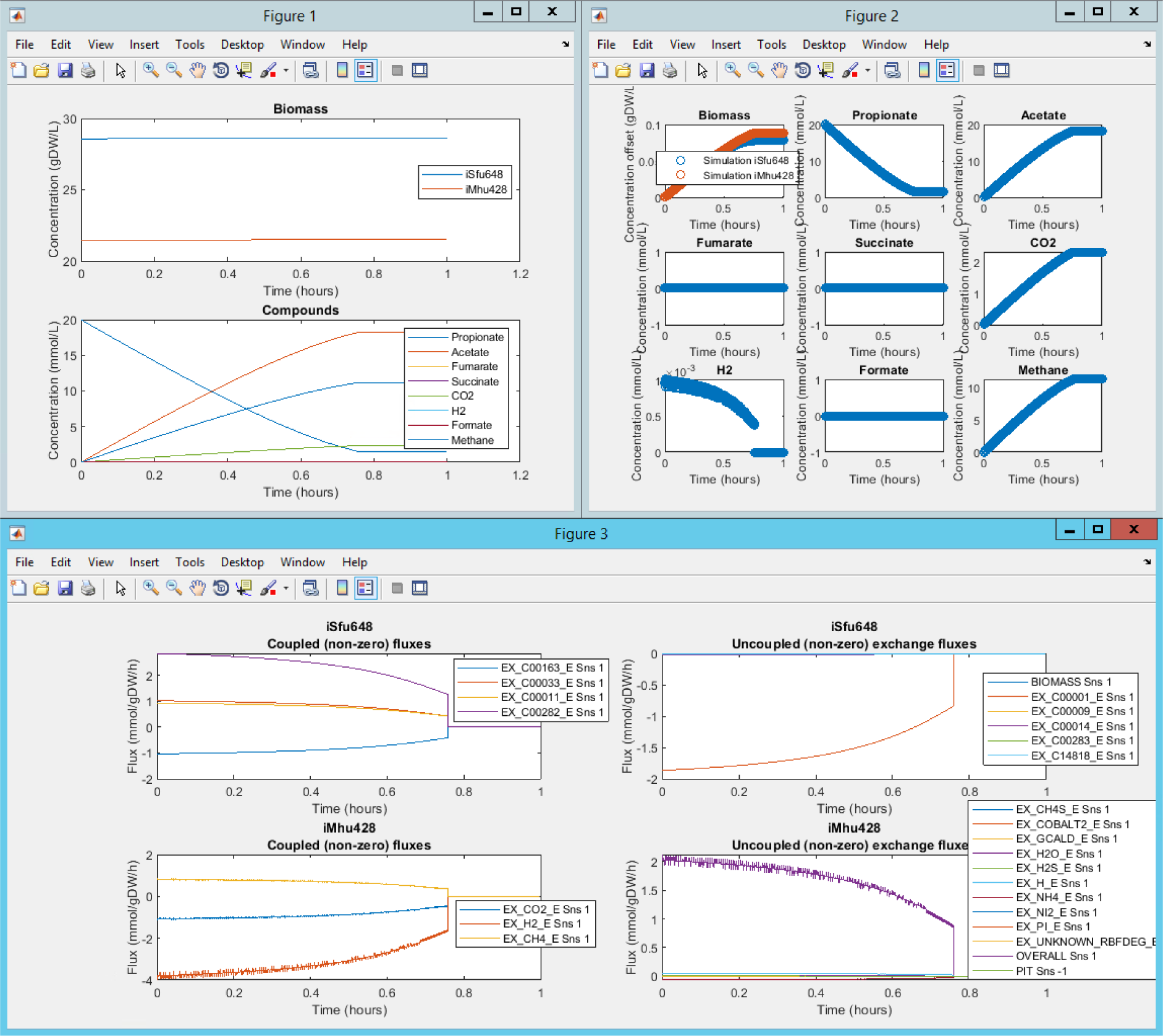
Automatically generated figure for Example 2. Simulation trajectory showing all microbiome member species and reactor compounds (Figure 1). Plot of biomass concentration as an offset to the initial concentration for all microbiome member species and individual plots for all reactor compounds (Figure 2). Plotting non-zero exchange fluxes over time which are coupled to reactor compounds (left), or not (right) for all microbiome member species (Figure 3).

#### Example 3: human gut microbiome with 773 species

Running Example 3 first requires the unpacking of the file `AGORA-1.01-Western-Diet.zip` containing the AGORA model collection. Note that for running the simulation with all species, 64GB of RAM are necessary (loading models as a Matlab data structure after the initial loading as SBML files cuts memory demand in half). The simulation of batch growth can then be started by `microbialSimMain(3)`. Simulation time can considerably be reduced by setting the parameter `solverPars.maxDeviation` to `"inf" in microbialSimMain.m` at the expense of numerical accuracy. Note that also arbitrary subsets of the model collection can be selected for the simulation (see commented example in the code).

**Fig S3.**
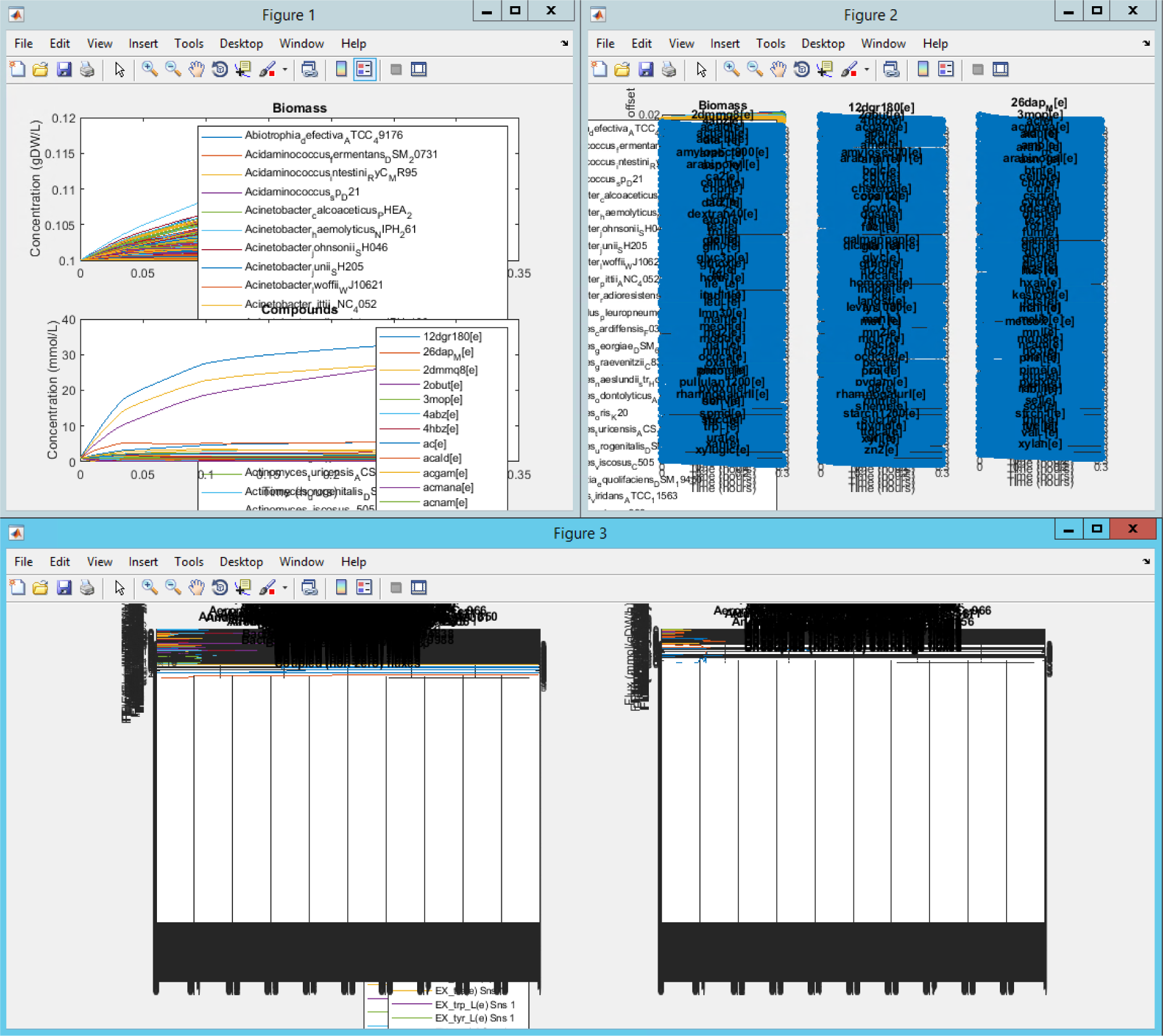
Automatically generated figure for Example 3. Simulation trajectory showing all microbiome member species and reactor compounds (Figure 1). Plot of biomass concentration as an offset to the initial concentration for all microbiome member species and individual plots for all reactor compounds (Figure 2). Plotting non-zero exchange fluxes over time which are coupled to reactor compounds (left), or not (right) for all microbiome member species (Figure 3).

